# Anticipatory Capture of Circulating Peptidergic Vesicles in a Clock Neuron

**DOI:** 10.1101/2025.10.22.684038

**Authors:** Markus K. Klose, Junghun Kim, Brigitte F. Schmidt, David L. Deitcher, Edwin S. Levitan

## Abstract

Neuropeptide release by *Drosophila* sLNv clock neurons controls circadian behaviors and sleep. Remarkably, neuropeptide content in sLNv terminals is rhythmic with late-night accumulation occurring while the axon arbor is expanding in preparation for midmorning synaptic exocytosis of neuropeptide-containing dense-core vesicles (DCVs). Past studies showed increased synaptic neuropeptide content can be produced by delivery of more neuropeptide to terminals or activity-dependent capture of circulating DCVs. To distinguish between these mechanisms, neuropeptide-containing DCVs were imaged in the living brain explant. First, post-exocytosis DCV axonal transport and synaptic neuropeptide accumulation following retrograde transport inhibition show that sLNv DCVs circulate. Furthermore, anterograde transport to terminals is constant throughout the day demonstrating there is no increase in DCV delivery. Rather, capture of circulating DCVs produces the daily boost in terminal neuropeptide content. Remarkably, this capture occurs before the daily increase in Ca^2+^ spike activity and is also independent of concurrent IP_3_ signaling and daily axon arbor expansion. Finally, a *per* clock gene mutation inhibits rhythmic DCV capture. Thus, rather than participating in axonal plasticity or responding to Ca^2+^ signaling, capture of circulating DCVs in sLNv presynapses is increased by the molecular clock in anticipation of activity-induced release hours later.

## Introduction

*Drosophila* sLNv clock neurons release two neuropeptides (PDF and sNPF) that are co-packaged in the same dense-core vesicles (DCVs) (Yu *et al*., 2025) to regulate circadian behavior (Crespo-Flores and Barber, 2022) and sleep (Shang *et al*., 2013). Interestingly, PDF content in sLNv neurons is rhythmic based on the function of the circadian clock (Park *et al*., 2000), which is a feature shared by other neuropeptides in other clock neurons (Hermann-Luibl *et al*., 2014; Fujiwara *et al*., 2018; Diaz *et al*., 2019; Sekiguchi *et al*., 2024). For both PDF and an exogenously expressed fluorescent protein-tagged neuropeptide, activity-dependent DCV exocytosis decreases presynaptic sLNv terminal neuropeptide content at midmorning (Park *et al*., 2000; Klose *et al*., 2021, 2025). Furthermore, this synaptic release is preceded by a late-night post-translational upregulation of terminal neuropeptide content (Park *et al*., 2000; Klose *et al*., 2021) that is advantageous because synaptic neuropeptide release scales with presynaptic neuropeptide content (Bulgari *et al*., 2014). However, the mechanism for the daily increase in terminal neuropeptide content is not known.

Traditionally, it was thought that newly synthesized DCVs are delivered to terminals from the soma for exocytotic release by anterograde axonal transport and eventually delivered back to the soma for degradation by retrograde axonal transport (Alonso and Assenmacher, 1983). In this context, the observed decrease in soma neuropeptide content preceding the increase in terminal neuropeptide content in sLNv terminals (Park *et al*., 2000; Klose *et al*., 2021) is consistent with a surge in anterograde transport producing the daily increase in terminal neuropeptide content (Fig. 1Ai). However, a recent study showed there is late-night DCV exocytosis at the sLNv soma (Klose *et al*., 2021), which might fully account for the somatic neuropeptide decrease. In the latter case, another mechanism must explain the daily neuropeptide increase in terminals. Such a mechanism was discovered in the peripheral nervous system where activity induces capture of DCVs as they circulate between distal terminal boutons and an axonal region near the soma to replenish synaptic neuropeptide content following stimulated neuropeptide release (Shakiryanova *et al*., 2006; Wong *et al*., 2012; Cavolo *et al*., 2016). Experimentally, induced capture is evident from a drop in retrograde DCV transport that accompanies increased synaptic neuropeptide content (Shakiryanova *et al*., 2006) (Fig. 1Aii). Thus, two mechanisms for increasing synaptic neuropeptide content are distinguished by their effects on axonal transport: either retrograde vesicle transport is decreased by synaptic capture of vesicles undergoing circulation or anterograde transport is increased to upregulate neuropeptide delivery to synapses. Although DCV axonal transport has been imaged in the living brain (e.g., Knabbe *et al*., 2018; Nassal *et al*., 2022), the role of these mechanisms for regulating daily changes in neuropeptide content is not known.

**Figure 1.**
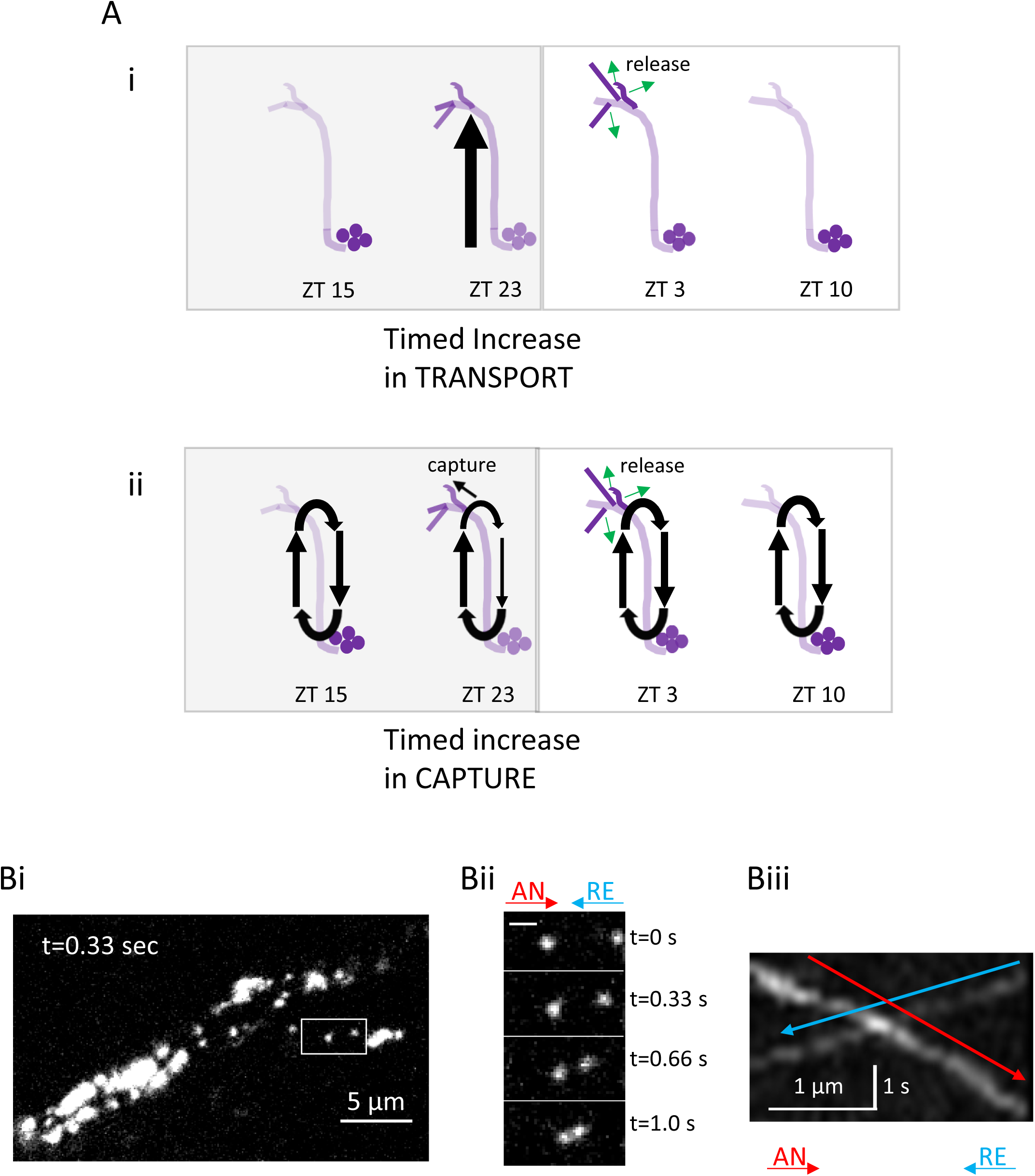
DCV transport in sLNv nerve terminal projections. A. Schematics present two hypotheses for how DCV transport dynamics could increase neuropeptide content at sLNv nerve terminals late at night and prior to release. Ai. Timed increase in anterograde transport of DCVs to nerve terminals late at night. Aii. Timed increase in capture of circulating DCVs late at night resulting in reduced retrograde transport. Bi. sLNv projections containing DCVs labeled with Dilp2-GFP were imaged at 3 Hz at ZT 3. White box reveals region of interest (ROI) for analysis. Bii. DCVs at t = 0, 0.33, 0.66, and 1.0 seconds (red arrow indicates anterograde (AN) transport and blue arrow indicates retrograde (RE) transport, scale bar = 1 µm). Biii. Kymograph of ROI (red arrow anterograde, blue arrow retrograde transport).

Here, axonal transport and presynaptic accumulation of neuropeptide-containing DCVs are imaged in the adult *Drosophila* brain. First, time-lapse imaging following synaptic kiss and run exocytosis and genetic inhibition of retrograde transport demonstrate DCVs in sLNv axons are circulating. Furthermore, rhythmic accumulation of DCVs in sLNv terminals occurs without a change in anterograde axonal transport. Instead, a late-night decrease in retrograde transport reveals an enhancement of vesicle capture that upregulates neuropeptide content in the sLNv terminals. Because DCV capture occurs hours before the daily rise in sLNv neuron activity (Klose *et al*., 2025), follow-up experiments address whether rhythmic presynaptic DCV capture depends on concurrent IP_3_ signaling (Klose *et al*., 2021), axon arbor plasticity (Fernandez et al., 2008; Ispizua et al., 2025) and the circadian clock.

## Results and Discussion

### DCV axonal transport is consistent with vesicle circulation in a clock neuron

To address the basis of the daily upregulation of neuropeptide content in sLNv terminals, we imaged DCV axonal transport in the adult brain explant from animals with cell specific expression of the DCV marker Dilp2-GFP (Wong *et al*., 2012), which recapitulates the known PDF content rhythm in sLNv terminals (Klose *et al*., 2021). Time-lapse imaging in the morning detected DCVs in distal dorsal sLNv axons undergoing both anterograde and retrograde axonal transport (Fig. 1B).

As capture of circulating DCVs could contribute to the daily neuropeptide rhythm in sLNv clock neurons, we sought to test for vesicle circulation in these brain neurons. Vesicle circulation was established by imaging the body wall innervation of the 2-dimensional filleted larva and conducting FRAP (fluorescence recovery after photobleaching) experiments with a scanning confocal microscope (Shakiryanova et al., 2006; Wong et al., 2012; Cavolo *et al*., 2016). These experiments are not feasible in the 3-dimensional brain explant because light scattering limits illumination and collection of fluorescence, and photobleaching illumination produces diffusing radicals that induce extensive photodamage. Therefore, we used an alternative approach based on imaging with the spinning disk confocal microscope, which is very sensitive and produces relatively little photobleaching. Specifically, experiments took advantage of (a) synaptic neuropeptide release being dominated by kiss and run DCV exocytosis (Wong *et al*., 2015; Bulgari *et al*., 2019, 2023; Klose *et al*., 2021), and (b) the fate of DCVs following exocytosis differing for the traditional model of DCV transport and vesicle circulation: with the former, post-exocytosis DCVs would not undergo anterograde transport because that process is reserved for newly synthesized vesicles, while with circulation, both anterograde and retrograde transport are expected.

Therefore, DCVs were selectively labeled by cell specific expression of a neuropeptide tagged with a fluorogen-activating protein (Dilp2-FAP) and application of the membrane impermeant fluorogen MG-TCarb (Bulgari *et al*., 2019; Klose *et al*., 2021). With this approach, fluorescence is only produced when the fluorogen diffuses through the fusion pore formed during kiss and run exocytosis to bind the FAP in the DCV lumen. Therefore, fluorescent DCVs were imaged in distal sLNv axons following the ∼1-hour incubation with MG-TCarb approaching the midmorning peak of synaptic DCV exocytosis and endogenous neuropeptide release at sLNv terminals (i.e., at ZT 2-3) (Klose *et al*., 2021, 2025). In animals co-expressing Dilp2-FAP and Dilp2-GFP, FAP and GFP colocalized in moving axonal puncta (Fig. 2A). Thus, as was found at the neuromuscular junction (NMJ), DCVs were FAP-labeled by kiss and run exocytosis (Bulgari *et al*., 2019). Furthermore, these post-exocytosis DCVs displayed both anterograde and retrograde axonal transport (Fig. 2 B,C), which is consistent with delivery of neuropeptides to terminals by vesicle circulation (Fig 1A ii).

**Figure 2.**
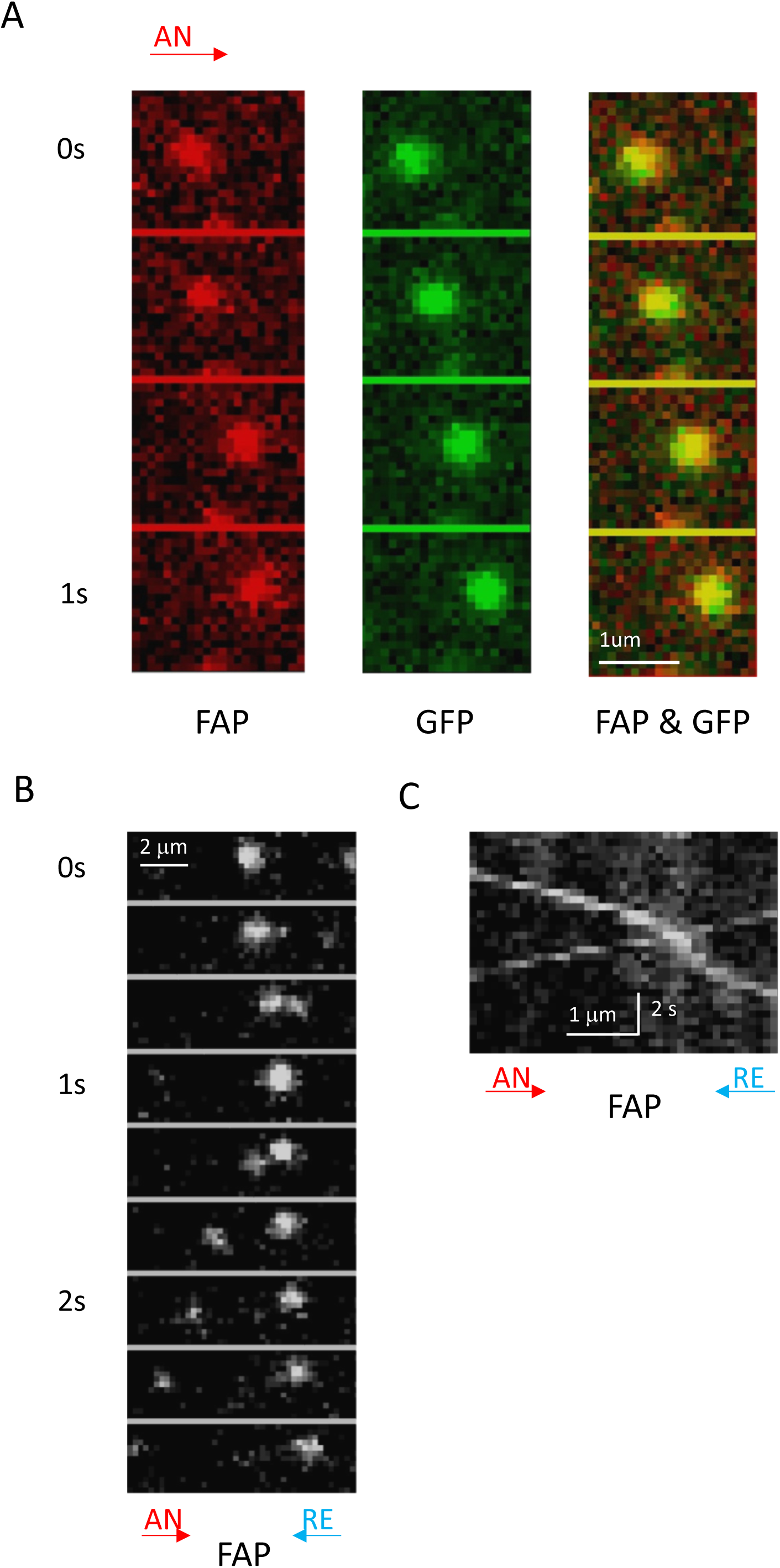
Post-exocytosis DCVs display anterograde and retrograde axonal transport. A. Co-localization of GFP and FAP signal in moving puncta through sLNv nerve terminals of *UAS-Dilp2-GFP, UAS-Dilp2-FAP; PDF-GAL4* flies (slope down, left to right). AN= anterograde transport (red arrow). B. FAP-labeled puncta in sLNv nerve terminals reveal both anterograde and retrograde mobility. RE= Retrograde transport (blue arrow). C. Kymograph revealing both anterograde (slope down, left to right) and retrograde (slope down, right to left) DCV axonal transport in sLNv projections.

With vesicle circulation through *en passant* boutons, retrograde transport out of the most distal bouton removes excess DCVs supplied by anterograde transport (Wong *et al*., 2012). Because dynein-mediated retrograde transport activation depends on the dynactin complex, this retrograde transport is inhibited by overexpressing the dynactin 2, p50 subunit (also known as dynamitin, dmn) (Burkhardt et al., 1997) resulting in accumulation of neuropeptide-containing DCVs in the most distal *en passant* bouton (Wong *et al*., 2012). Therefore, to independently test if DCVs undergo vesicle circulation in sLNv terminals, *UAS-dmn* was used to overexpress the dynactin subunit in *PDF>Dilp2-GFP* animals. Figure 3 shows that the neuropeptide content of the most distal sLNv boutons in *PDF>Dilp2-GFP* animals examined during either the first day or the fifth day following eclosion are not different than their proximal neighboring boutons (CON #1 vs. CON #2, Figure 3Bi and ii). However, dmn overexpression to inhibit retrograde transport resulted in ∼10-fold more neuropeptide accumulation in the most distal bouton at both time points (dmn #1 vs. dmn #2, Figure 3Bi and ii). Thus, a genetic perturbation verifies DCVs in sLNv terminals are circulating.

**Figure 3.**
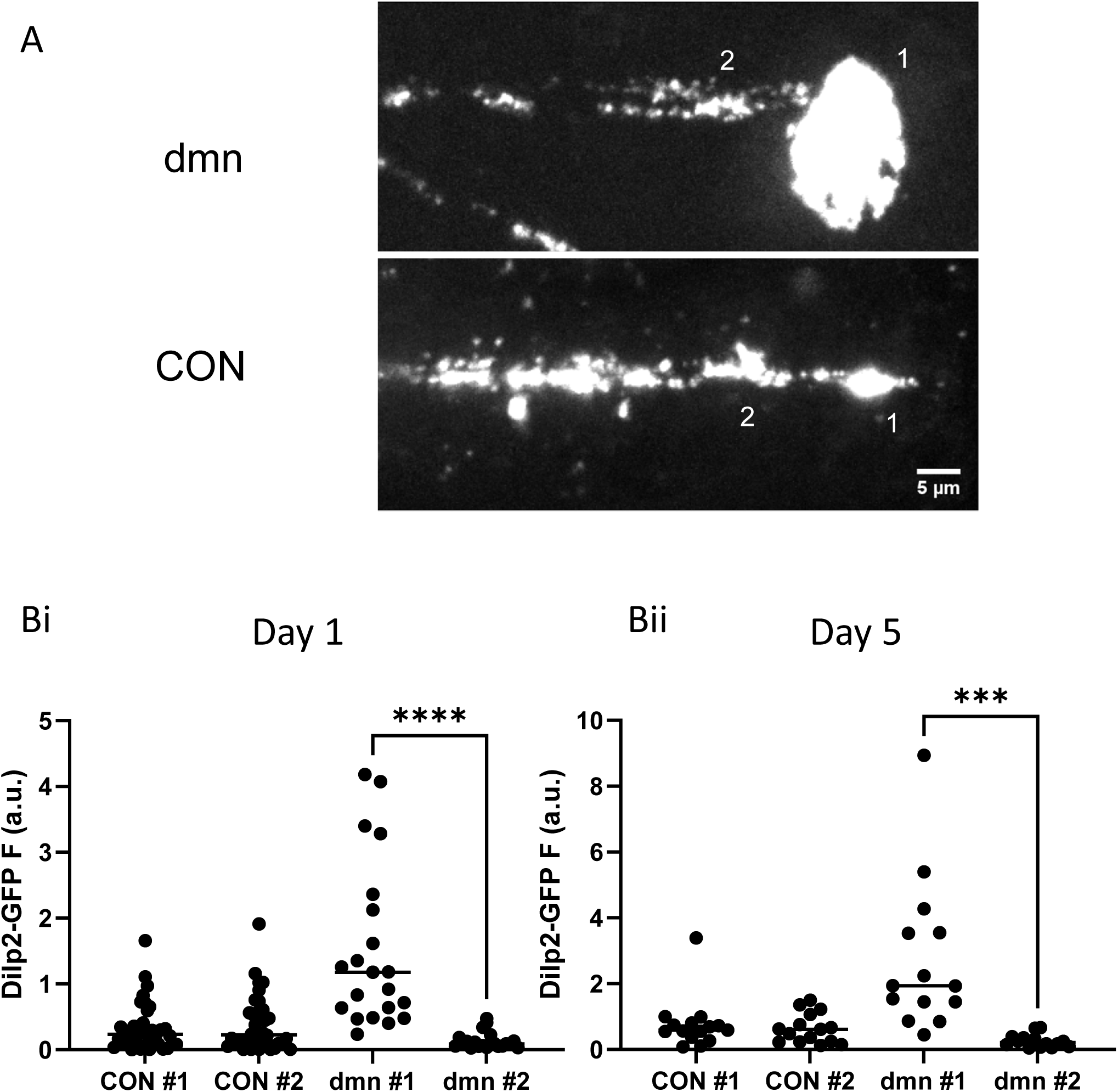
Dynamitin increases neuropeptide content of sLNv distal terminal boutons. A. Images from distal sLNv nerve terminals of *UAS-dmn/UAS-Dilp2-GFP; PDF-GAL4* flies (dmn) and *UAS-Dilp2-GFP; PDF-GAL4* control flies (CON) at ∼ZT 2 on Day 1 after eclosion. B. Dilp2-GFP fluorescence was compared between the most distal *en passant* bouton (#1) and its proximal neighbor bouton (#2) revealing a dramatic content difference between the two not see in control terminals at the first (i) and fifth (ii) day post-eclosion. ***P < 0.001, ****P < 0.0001, Paired t-tests with Welch’s correction for different variances. Points represent data from individual neurons (N = 20-36 from 7-10 hemibrains in i, and 14-15 from 5-6 hemibrains in ii).

### Rhythmic capture of circulating DCVs in sLNv terminals

To test for regulated axonal transport of DCVs, we initially measured the number of vesicles being transported per minute (i.e., flux) and calculated the ratio of anterograde to retrograde DCV flux (A/R) (Fig. 4A,B) showed that the flux ratio changed (P < 0.001, one-way ANOVA) with a peak at ZT 23 (Fig. 4B). Therefore, there is a net increase in axonal transport of DCVs to sLNv terminals late at night in preparation for the midmorning burst of synaptic neuropeptide release.

**Figure 4.**
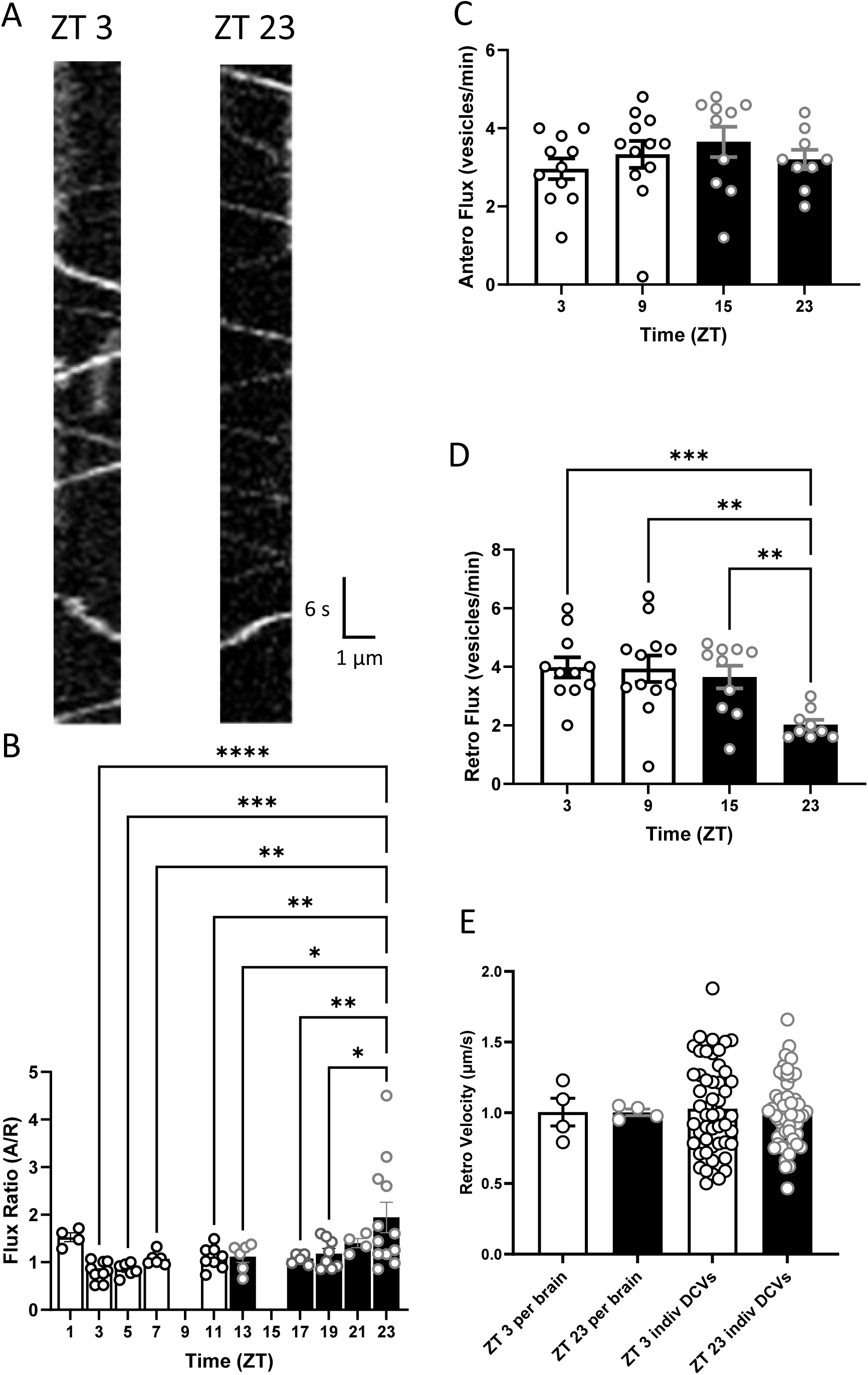
DCV flux and velocity in sLNv nerve terminals show late-night capture upregulates synaptic neuropeptide content. A. Kymographs reveal both anterograde (slope down, left to right) and retrograde (slope down, right to left) DCV axonal transport in sLNv projections of *PDF>Dilp2-GFP* flies at ZT 3 and ZT 23. B. Flux ratios (anterograde/retrograde) at different times of the day. n values (number of hemibrains) for ZT 1, 3, 5, 7, 11, 13, 17, 19, 21, 23 are 4, 10, 6, 6, 8, 6, 8, 8, 4, 12, respectively. One-way ANOVA revealed significant difference (P < 0.001). Post-test analysis by Dunnett’s multiple-comparison test is presented: ****P < 0.0001, ***P < 0.001, **P < 0.01, *P < 0.05. C. Anterograde flux and D. Retrograde flux (vesicles/min) in *PDF>Dilp2-GFP* brains. For D, the n values for ZT 2-4, 8-10, 14-16, and 23 are 11, 12, 10 and 9, respectively. P < 0.01, Brown-Forsythe ANOVA. ***P < 0.001, **P < 0.01, Dunnett’s T3 multiple comparisons test. E. Retrograde velocity of DCVs at ZT 3 and 23 in *PDF>Dilp2-GFP* brains (left, average per brain; ZT 3 n = 4, ZT 23 n = 4; right, individual vesicle measurements: ZT 3 n = 49, ZT 23 n = 51).

Anterograde and retrograde DCV transport fluxes were then examined to determine which component of the flux ratio changed, the numerator (A) or the denominator (R). Analysis of axonal transport showed that anterograde DCV flux (A) was constant at all times tested (Fig. 4C), thus ruling out rhythmic changes in the ratio being driven by changes in anterograde axonal transport from the soma. Rather, retrograde axonal DCV flux (R) cycled (P < 0.01, one-way ANOVA) with a minimum late at night at ZT 23 (Fig. 4D). This drop in retrograde flux cannot be explained by exocytosis-induced DCV depletion since the flux drop precedes daily synaptic DCV exocytosis and native synaptic neuropeptide release by several hours (Klose *et al*., 2021, 2025). Furthermore, the decrease in retrograde flux occurred without altering the velocity of DCVs undergoing retrograde axonal transport (Fig. 4E), thereby excluding a slowing of the retrograde transport motors. The observed reduction in retrograde flux hours before synaptic release with no change in retrograde velocity is indicative of increased synaptic capture of circulating DCVs (Bulgari *et al*., 2006; Wong *et al*., 2012; Cavolo *et al*., 2016). Therefore, increased DCV capture, as opposed to greater DCV delivery by anterograde transport, produces the daily increase in synaptic neuropeptide content.

### DCV capture is independent of rhythmic activity, IP_3_ signaling and axon plasticity

The late-night (ZT 23) synaptic capture of DCVs precedes the daily rise in Ca^2+^ spiking, which peaks 4 hours later at midmorning (Klose *et al*., 2025). However, Ca^2+^ release from endoplasmic reticulum participates in activity-dependent DCV capture at the NMJ (Wong *et al*., 2009) and there is IP_3_-IP3R signaling in sLNv neurons at ZT 23 (Klose *et al*., 2021). Therefore, we tested for a role of IP_3_ signaling in late-night synaptic DCV capture. Specifically, the flux ratio at ZT 23 was determined after cell specific expression of IP_3_ sponge (Uchiyama *et al*., 2002), which sequesters IP_3_. IP_3_ sponge did not block capture at ZT 23 (Fig. 5A). Therefore, Ca^2+^ elevation induced by either activity or IP_3_ signaling is not required for late-night DCV capture in the presynapse.

**Figure 5.**
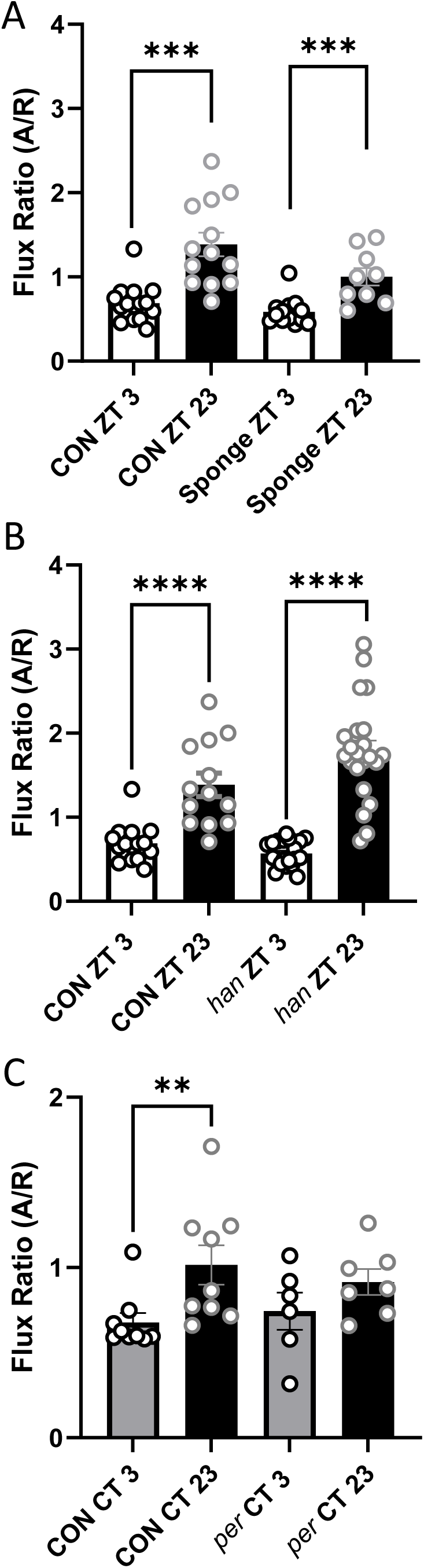
Rhythmic DCV capture requires the clock gene *per*, but not concurrent IP_3_ signaling or PDFR-dependent axonal plasticity. A. Ratios of anterograde to retrograde DCV flux (A/R) at ZT 3 and ZT 23 in *w*; *UAS-Dilp2-GFP; PDF-GAL4* controls (CON) and IP_3_ sponge; *UAS-Dilp2-GFP; PDF-GAL4 UAS-IP_3_ sponge* sLNv axons. n ≥ 13 for CON; n > 9 for IP_3_ sponge. For both CON and *han*. B. Ratios of anterograde to retrograde DCV flux (A/R) at ZT 3 and ZT 23 in *w*; *UAS-Dilp2-GFP; PDF-GAL4* controls (CON) and *han*; *UAS-Dilp2-GFP; PDF-GAL4* PDFR null (*han*) sLNv axons. n ≥ 13 for CON; n > 20 for *han*. For both CON and *han*. C. Ratios of anterograde to retrograde DCV flux (A/R) 2 days after transition to constant darkness at CT3 and CT23 in *w*; *UAS-Dilp2-GFP; PDF-GAL4* controls (CON) and *y^1^per^01^w\**; *UAS-Dilp2-GFP; PDF-GAL4* (*per*) sLNv axons. n = 9 for CON; n = 6 for CT3 and n = 7 for CT23 in *per* animals. All data in this figure were analyzed using the Mann-Whitney test. **P < 0.01, ***P< 0.001, ****P < 0.0001.

We then considered whether the rhythmic DCV capture is linked to daily morphological plasticity in the sLNv axonal arbor (Fernandez *et al*., 2008), which is associated with varicosities (i.e., boutons) containing more DCVs (Ispizua et al., 2025). For this purpose, DCV axonal transport was studied in a PDF receptor (PDFR) null mutant (*han*), which prevents daily changes in terminal morphology while preserving rhythmic changes in PDF (Herrero *et al*., 2020). In PDFR null animals, the daily rhythmic changes in axonal flux ratio persisted (Fig. 5B). Therefore, the daily rhythm in DCV capture occurs independently of PDFR expression and the accompanying axonal plasticity it drives.

### Rhythmic DCV capture requires the circadian clock

The PDFR null mutant also abolishes morning anticipation behavior (i.e., the increase in motor activity prior to sunrise) (Hyun *et al*., 2005). Because rhythmic capture occurs concurrently with this clock-dependent increase in locomotor behavior but persists when morning anticipation is disrupted by the PDFR null mutation, we considered whether capture depends on the circadian clock. If capture is independent of the circadian clock, rhythmic capture would likely be driven through environmental cues, namely daily changes in light levels, and thus would be disrupted after shifting to a constant level of illumination. However, after shifting to constant darkness (DD) capture remained different between CT 23 and CT 3 (Fig. 5C, CON). To further test for a role of the circadian clock under DD conditions, we examined the effect of a null mutation in the *period* gene (*per^01^*), which is a critical component of the molecular clock. Consistent with the loss of oscillation in synaptic PDF content with this mutant (Park *et al*., 2000), elevated capture under DD conditions was disrupted in *per^01^* animals (Fig. 5C, *per*). Therefore, the circadian clock controls the rhythmic increase in the presynaptic neuropeptide pool by vesicle capture.

### Conclusions

In *Drosophila* sLNv clock neurons, there is a daily post-translational surge in synaptic PDF content (Park *et al*., 2000), which is also evident with an exogenous neuropeptide (Klose *et al*., 2021). Here, imaging neuropeptide-containing vesicles in the *Drosophila* brain explant, along with genetic perturbations of dynactin and *per*, demonstrates clock-dependent capture of circulating DCVs elevates presynaptic neuropeptide content in sLNv neurons each day. Because synaptic neuropeptide release is proportional to presynaptic neuropeptide content (Bulgari *et al*., 2014), midmorning peptidergic synaptic transmission that controls circadian behavior via PDF is promoted by clock-dependent late-night synaptic capture of circulating DCVs.

Clock-dependent vesicle capture likely also upregulates presynaptic sNPF, a neuropeptide that is co-packaged in the same sLNv neuron DCVs as PDF and promotes nighttime sleep (Shang *et al*., 2013; Yu *et al*., 2025). Unlike PDF, sNPF gene expression in sLNv neurons is rhythmic (Kula-Eversole *et al*., 2010; Ma *et al*., 2021), suggesting that both neuropeptide synthesis in the soma and vesicle capture in terminals act synergistically to rhythmically upregulate presynaptic sNPF stores for peptidergic transmission. It will be of interest to determine how clock neurons in *Drosophila* and mammals utilize clock-regulated vesicle capture and changes in neuropeptide gene expression to control rhythmic behaviors.

Two mechanisms of DCV capture were discovered in the fly NMJ, constitutive capture, which maintains neuropeptide content during quiescence by sequestering DCVs undergoing bidirectional transport, and activity-dependent capture, which replenishes presynaptic neuropeptide stores for minutes following release by tapping into anterograde DCV flux (Shakiryanova et al., 2006; Wong *et al*., 2012; Cavolo *et al*., 2016). In contrast to activity-dependent capture, capture in sLNv clock terminals is not induced by activity, but rather is driven by the clock (Fig. 5) hours in advance of future synaptic activity and release. This predictive DCV capture is independent of sLNv axonal plasticity (Fig. 4C), thus excluding daily neuropeptide accumulation simply reflects capture to occupy the expanding axonal arbor with a greater number of boutons. Also, daily DCV capture does not require IP_3_ signaling, which is concurrent with, but independent of, increased capture (Fig. 4A,B). A future challenge will be to resolve how clock-induced vesicle capture operates compared to activity-dependent and constitutive capture mechanisms, which are not fully understood in *Drosophila*.

## Materials and Methods

### Imaging

Flies were entrained for at least for 72 hours in a 12-hour light: 12-hour dark (LD) schedule before dissection of 4-9 day old males to generate brain explants. Dissections during the dark phase were performed under a red light. Adult flies were immobilized with CO_2_ gas and brains were dissected in 0 Ca^2+^ HL3 saline solution (70 mM NaCl, 5 mM KCl, 20 mM MgCl_2_*6H_2_O, 115 mM Sucrose, 5 mM Trehalose, 5 mM Hepes, and 10 mM NaHCO_3_, pH 7.3) and brain explants were then put into polylysine-coated plastic dishes containing HL3 with 2 mM Ca^2+^ (Klose et al., 2016). sLNv somas are located on the ventral side of the Drosophila brain and send axons to the dorsal protocerebrum. Therefore, brain explants were viewed from the dorsal surface of the brain so that DCVs in the distal axon could be imaged with the dipping water immersion objective of an upright spinning disk confocal microscope. Imaging was done on setups with an upright Olympus microscope equipped with a 60× 1.1 NA dipping water immersion objective, a Yokogawa spinning disk confocal head, a Teledyne Photometrics sCMOS camera, and lasers for illumination (488 nm for GFP and 640 nm for FAP). In some experiments, a recombinant with UAS-Dilp2-GFP and UAS-Dilp2-FAP was used so that GFP could be viewed for focusing before application of the membrane impermeant fluorogen MG-Tcarb (Klose *et al*., 2021). For DCV axonal transport, two axonal regions in the dorsal protocerebrum were imaged in each hemisphere of the brain. Time-lapse experiments entailed acquiring 180 images with 35-150 ms exposure times at 3 Hz. From these movies, regions where dorsally projecting axons defasciculated from bundles were used for generating kymographs with the Kymograph plugin of ImageJ. Quantification was done in Imagej or Fiji. Statistical analysis (i.e., tests and calculation of standard error of the mean (SEM) for error bars) was performed with Graphpad Prism software.

### Fly lines

All flies used the *PDF*-*GAL4* promoter on the third chromosome (provided by Paul Taghert, Washington University in St. Louis). *UAS-Dilp2-GFP* and *UAS-Dilp2-FAP* were reported previously (Wong *et al*., 2012; Bulgari *et al*., 2019). *w^1118^*flies were from Zachary Freyberg (University of Pittsburgh). Fly lines from the Bloomington *Drosophila* stock center included *Bl#* 33068 (*han*) and *Bl#* 80917 (*per^01^*). The IP_3_ sponge line was provided by C. Andrew Frank (University of Iowa).

## Acknowledgments

We thank Drs. C.A. Frank for kindly providing IP_3_ sponge flies. Stocks obtained from the Bloomington Drosophila Stock Center (NIH P40OD018537) were used in this study. Research reported in this publication was supported by the National Institute Of Neurological Disorders And Stroke of the National Institutes of Health under Award Number R01NS032385 to ESL. The content is solely the responsibility of the authors and does not necessarily represent the official views of the National Institutes of Health.

## Notes

The authors declare no competing interests.

### Competing Interest Statement

The authors have declared no competing interest.

### Summary of Updates

New data in Figure 3 shows the effect dynamitin overexpression on neuropeptide content in sLNv nerve terminals supporting the conclusion that circulating DCVs are captured into nerve terminals to increase content for subsequent release.

